# MLL3/4 methyltransferases regulate the differentiation of pluripotent stem cells through coordinating glycolysis and mitochondrial respiration

**DOI:** 10.64898/2026.03.24.713976

**Authors:** Suza Mohammad Nur, Yunbo Jia, Muyi Ye, Caylin A. Lepak, Issam Ben-Sahra, Kaixiang Cao

## Abstract

Enhancer-regulating epigenetic modifiers play critical roles in normal physiological processes and human pathogenesis. The major enhancer regulator paralogs MLL3 and MLL4 (MLL3/4) belong to the lysine methyltransferase 2 (KMT2) family, which catalyzes the methylation of lysine 4 on histone H3 (H3K4me). MLL3/4 are required for enhancer activation and are essential for mammalian development and stem cell differentiation. Although recent studies have linked MLL3/4 with different metabolic pathways in the regulation of stem cell self-renewal and cancer cell growth, the mechanisms connecting enhancer function to metabolic control remain elusive. Here, using respiration flux assays, stable isotope tracing, transcriptomics, and stem cell biology techniques, we show that the loss of MLL3/4 impairs glycolysis and mitochondrial respiration in murine embryonic stem cells. Mechanistically, MLL3/4 deficiency suppresses the expression of the rate-limiting glycolytic enzyme hexokinase 2 (HK2) and compromises the function of the oxoglutarate dehydrogenase (OGDH) complex, thereby coordinately impairing central carbon metabolism. Remarkably, combined restoration of HK2 and OGDH rescues the metabolic defects caused by MLL3/4 loss and reinstates differentiation capacity. Taken together, our study identifies a direct link between enhancer-regulating epigenetic machineries and metabolic control of cell fate transition, providing a mechanistic framework for understanding how enhancer malfunction contributes to developmental abnormalities and human diseases.

## Introduction

Enhancer deregulation is a major driver of human congenital disorders (*1-4*). In mammals, nucleosomes at enhancers are mono-methylated on lysine 4 of histone H3 (H3K4) by the paralogous enzymes Mixed-lineage Leukemia 3 (MLL3, also known as KMT2C) and Mixed-lineage Leukemia 4 (MLL4, also known as KMT2D) (*5-9*). Mice null for MLL4 die shortly after gastrulation, while MLL3 loss leads to neonatal lethality (*10, 11*). Heterozygous loss-of-function mutations in MLL4 lead to Kabuki syndrome (*1*) and those in MLL3 cause Kleefstra syndrome 2 (*4, 12, 13*). Both disorders are characterized by craniofacial dysmorphism and developmental delay, highlighting the instructive role of MLL3/4 in normal development and human pathogenesis. However, molecular mechanisms by which MLL3/4 function in biologically normal and pathological processes remain poorly understood.

Recent results have revealed a functional link between MLL3/4 and cellular metabolism. MLL3/4 and nucleotide synthesis pathways co-regulate the self-renewal of pluripotent stem cells (PSCs) (*14*). Moreover, steady-state abundance of glycolytic metabolites such as glucose-6-phosphate (G6P) and fructose-6-phosphate (F6P), and tricarboxylic acid (TCA) cycle metabolites including succinate, fumarate, and malate, is reduced in PSCs lacking MLL3/4 (*14*). In contrast, MLL4-deficient lung cancer and melanoma cells adopt upregulated glycolysis (*15, 16*), suggesting a context-dependent role of MLL3/4 in regulating cellular metabolism. How MLL3/4 modulate metabolic pathways and whether they govern cell fate via these pathways are largely unknown.

Glycolysis and mitochondrial respiration are tightly regulated in PSCs. Naive PSCs, such as murine embryonic stem cells (ESCs), exhibit metabolic bivalency, utilizing both glycolysis and mitochondrial respiration for energy production, whereas primed PSCs are predominantly glycolytic (*17*). On the other hand, somatic cells are more dependent on mitochondrial respiration (*18, 19*). Inhibition of glycolysis drives the exit from pluripotency, while stimulation of glycolysis enhances somatic cell reprogramming to induced PSCs (iPSCs) (*18, 20*), suggesting that the balance between glycolysis and mitochondrial respiration instructs cell fate transition. To date, how metabolic states are established during development and whether they instruct cell identity remain enigmatic.

To investigate the functional relationship of MLL3/4, cellular metabolism, and PSC differentiation, we examined the impact of MLL3/4 loss on glycolysis and mitochondrial respiration in ESCs. Both extracellular acidification rate (ECAR) and oxygen consumption rate (OCR) are reduced in MLL3/4-null ESCs. Stable isotope tracing reveals a reduction in the labeling of specific metabolites in both glycolysis and the TCA cycle upon MLL3/4 loss. MLL3/4 are required for the expression of hexokinase 2 (HK2) and for proper lipoylation of dihydrolipoamide S-succinyltransferase (DLST), the E2 subunit of the oxoglutarate dehydrogenase complex (OGDHc). Moreover, combined overexpression of HK2 and OGDH partially rescues the metabolic and differentiation defects caused by MLL3/4 loss. Our results provide novel insight into understanding the crosstalk between epigenetic machineries and cellular metabolism. Importantly, they may pave the way for developing novel strategies to target human diseases driven by enhancer deregulation.

## Results

### MLL3/4 loss dampens glycolytic activity in ground-state ESCs

We previously found that the loss of MLL3/4 in ESCs reduced the steady-state abundance of glycolytic intermediates, including G6P and F6P (*14*), suggesting that MLL3/4 regulate glycolysis in naive PSCs. To directly test this idea, we measured the ECAR of wildtype (WT) and MLL3/4 double knockout (DKO) ESCs (*14*) using a glycolysis stress assay. MLL3/4 deficiency markedly reduced both basal glycolysis and glycolytic capacity (Fig. 1A-C), suggesting that MLL3/4 are required to maintain the glycolytic activity in naive PSCs. To determine if the decreased steady-state abundance of glycolytic metabolites reflects impaired glucose metabolism, we next performed stable isotope tracing with [^13^C_6_]-glucose in WT and MLL3/4 DKO ESCs, followed by liquid chromatography-tandem mass spectrometry (LC-MS/MS) analysis (Fig. 1D). Consistent with results in steady-state metabolomics, the incorporation of ^13^C into F6P/G1P (glucose-1-phosphate)/G6P (m+6) was significantly reduced by MLL3/4 loss (Fig. 1E and Fig. S1A). Moreover, MLL3/4 loss reduced labeling of lactate (m+3) from [^13^C_6_]-glucose (Fig. 1F and Fig. S1B), corresponding to the decreased glycolytic activity in MLL3/4 DKO ESCs. To determine whether this defect was caused by reduced glucose import, we measured glucose uptake and found no significant difference between WT and MLL3/4 DKO ESCs (Fig. 1G). These findings indicate that MLL3/4 loss does not impair glucose entry into cells but instead disrupts early rate-limiting steps in glycolysis, thereby suppressing glycolysis in ESCs.

**Figure 1.**
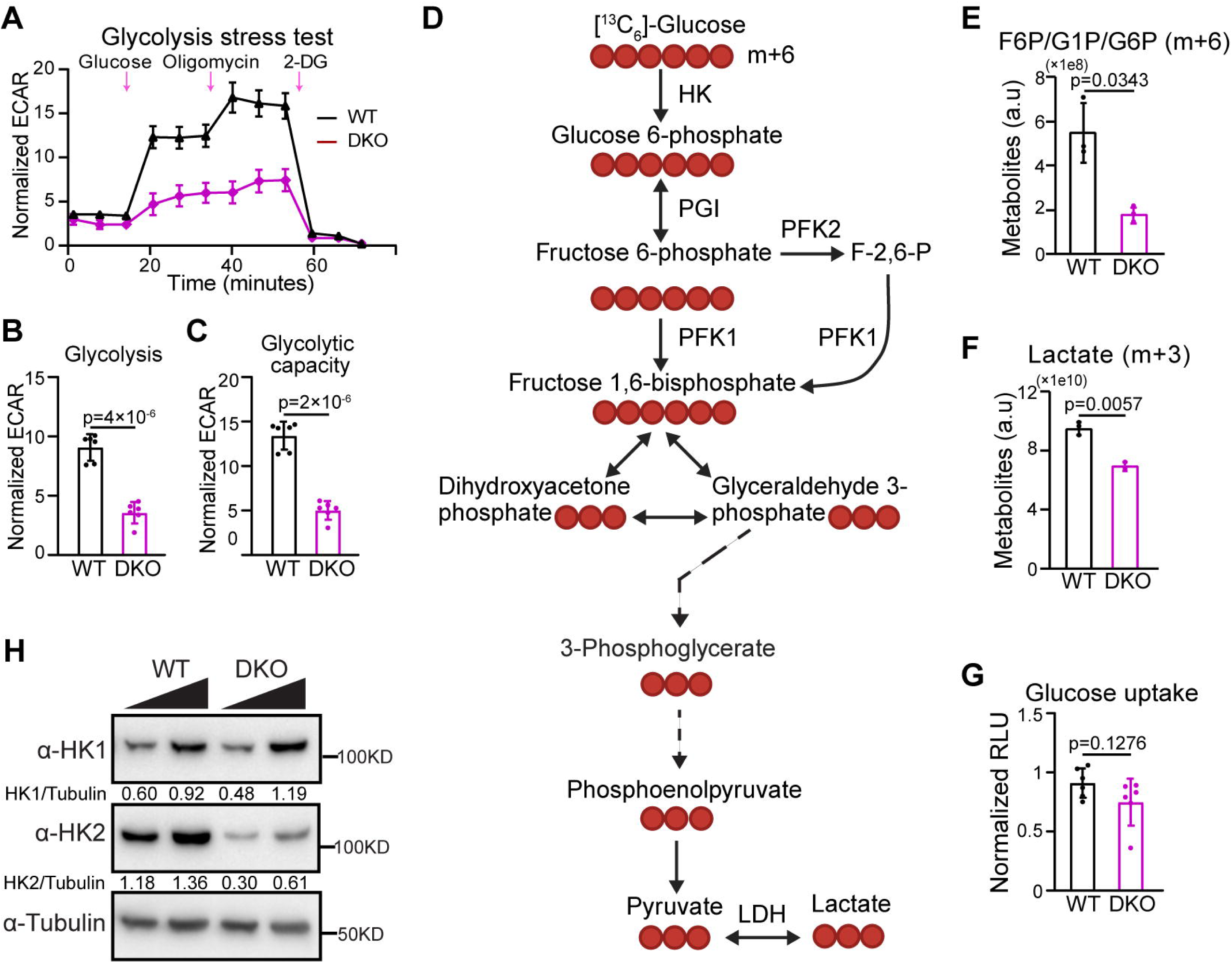
The loss of MLL3/4 leads to impaired glycolytic activity in ground-state ESCs. (A) Representative ECAR traces of WT and MLL3/4 DKO ESCs measured by the Seahorse XF glycolysis stress test assay. Data were normalized by the number of cells at the time of the assay and are represented as mean ± S.D. Each data point had six replicates, and the experiment was repeated three times independently. 2-DG: 2-Deoxy-D-glucose. (B-C) Bar plots of ECAR showing the levels of glycolysis (B) and glycolytic capacity (C) in WT and MLL3/4 DKO ESCs. Data are presented as mean ± S.D. p-values were calculated using Student’s t-test. (D) Schematic of glycolysis illustrating the fate of ^13^C carbons upon the incubation with [^13^C_6_]-glucose. (E-F) Bar plots showing the levels of F6P/G1P/G6P (m+6, E) and lactate (m+3, F) in WT and MLL3/4 DKO ESCs labeled with [^13^C_6_]-glucose. n=3 biological replicates per sample. Data are presented as mean ± S.D. p-values were calculated using Student’s t-test. (G) Bar plots showing the levels of glucose uptake in WT and MLL3/4 DKO ESCs. n=6 replicates per sample. Experiments were repeated three times independently and representative results are shown. Data were normalized by the number of cells at the time of the assay and are presented as mean ± S.D. p-values were calculated using Student’s t-test. RLU: relative light unit. (H) Western blots showing the levels of HK1 and HK2 in WT and MLL3/4 DKO ESCs. 10 and 20 μg of cell lysates were loaded for each sample. Experiments were repeated three times independently and representative results are shown. Protein levels in each lane were quantified by normalizing with Tubulin signals of the corresponding lane.

To elucidate the mechanism underlying this defect, we examined RNA-seq data (*14*) for genes encoding glycolytic enzymes involved in the production and utilization of G6P and F6P. Among these, *Hk2*, which encodes hexokinase 2, the enzyme that catalyzes the first committed step of glycolysis by converting glucose to G6P, was significantly downregulated in MLL3/4 DKO ESCs (Fig. S1C). On the other hand, the expression levels of genes encoding glucose-6-phosphate isomerase (*Gpi1*), phosphofructokinase 1 (*Pfkl* and *Pfkp*), and phosphofructokinase 2 (*Pfkfb1* and *Pfkfb2*) were largely unchanged (Fig. S1C). Consistent with the transcriptomic data, western blotting analysis showed that HK2 protein levels were markedly reduced by MLL3/4 loss, whereas levels of HK1 (hexokinase 1), the paralog of HK2, remained largely unaffected (Fig. 1H). To investigate how MLL3/4 regulate *Hk2* expression, we analyzed ChIP-seq datasets (*7, 14*) and found at least six putative enhancers near the *Hk2* locus, which were bound by MLL4 and exhibited reduced H3K27ac levels in MLL3/4 DKO ESCs (Fig. S1D). These data suggest that MLL3/4 could promote *Hk2* expression through these putative enhancers. Taken together, our results indicate that glycolysis in naive PSCs requires MLL3/4 and that the decreased HK2 levels may be responsible for the defective glycolysis caused by MLL3/4 loss.

### MLL3/4 are required for mitochondrial respiration in ground-state ESCs

Metabolic bivalency, defined by the concurrent use of glycolysis and mitochondrial respiration, is one of the hallmarks of naive PSCs (*21, 22*). To examine the role of MLL3/4 in regulating mitochondrial respiration in naive PSCs, we measured the OCR of WT and MLL3/4 DKO ESCs using a mitochondrial stress assay. Interestingly, MLL3/4 deficiency significantly reduced both basal and maximal respiration (Fig. 2A-C), suggesting that MLL3/4 are required to maintain the bivalent metabolic program of naive PSCs. To define where mitochondrial metabolism was disrupted, we performed stable isotope tracing with [^13^C_6_]-glucose labeling (Fig. 2D). MLL3/4 loss did not have a significant impact on the labeling of citrate/isocitrate (m+2) and alpha-ketoglutarate (α-KG; m+2) (Fig. 2E-F and Fig. S2A-B). In contrast, labeling of succinate (m+2), fumarate (m+2), and malate (m+2) was significantly reduced in MLL3/4 DKO ESCs (Fig. 2G-I and Fig. S2C-E). These findings suggest that MLL3/4 loss disrupts TCA cycle activity downstream of α-KG, thereby contributing to the impaired mitochondrial respiration.

**Figure 2.**
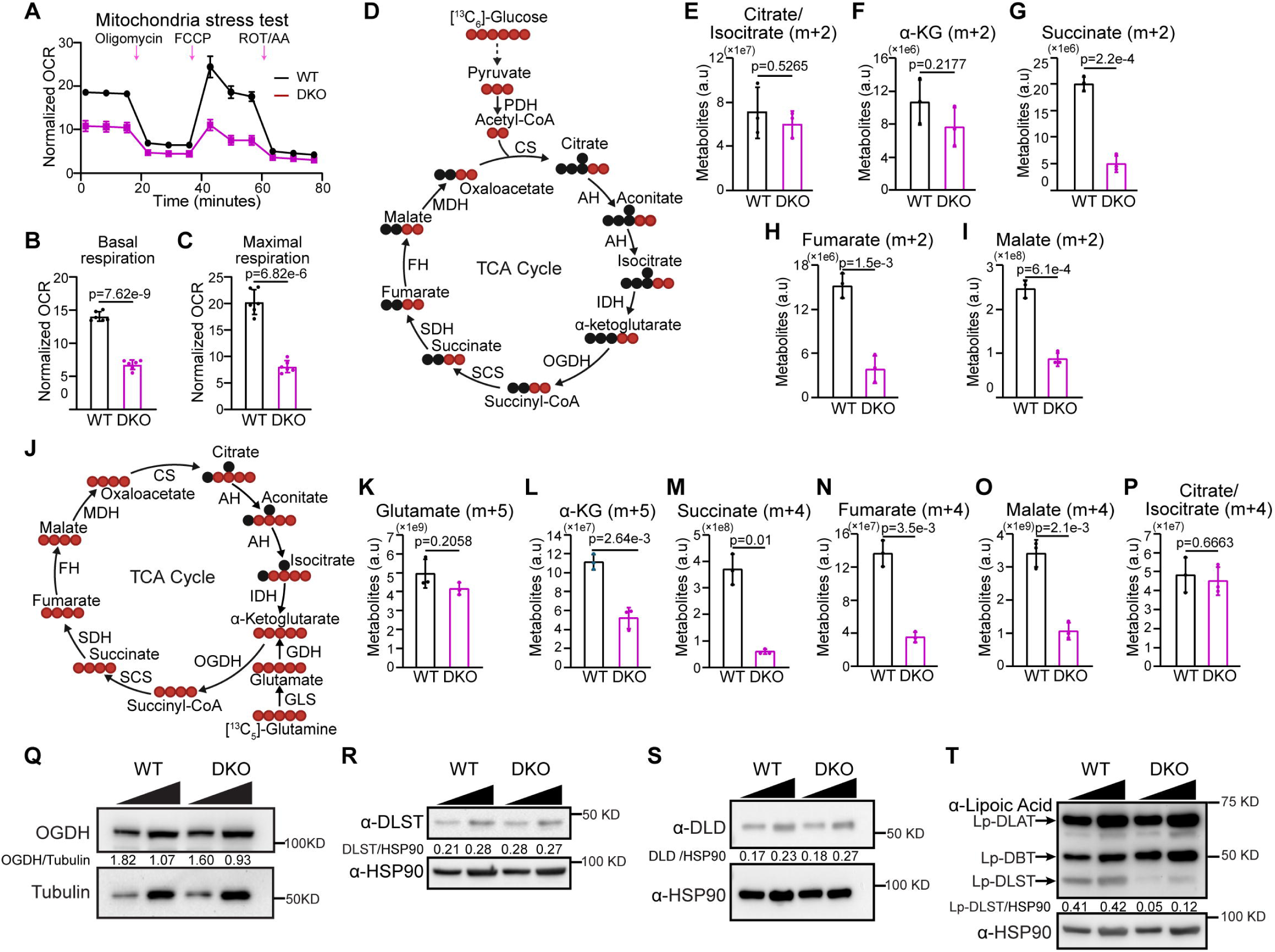
The loss of MLL3/4 leads to impaired mitochondrial respiration in ground-state ESCs. (A) Representative OCR traces of WT and MLL3/4 DKO ESCs measured by the Seahorse XF mitochondrial stress test assay. Data were normalized by the number of cells at the time of the assay and are represented as mean ± S.D. Each data point had six replicates, and the experiment was repeated three times independently. FCCP: Carbonyl Cyanide-4-(trifluoromethoxy) phenylhydrazone. ROT/AA: Rotenone and Antimycin A. (B-C) Bar plots showing the levels of OCR during basal (B) and maximal (C) respiration in WT and MLL3/4 DKO ESCs. Data are presented as mean ± S.D. p-values were calculated using Student’s t-test. (D) Schematic of the TCA cycle illustrating the fate of ^13^C carbons upon the incubation with [^13^C_6_]-glucose. (E-I) Bar plots showing levels of citrate/isocitrate (m+2, E), α-KG (m+2, F), succinate (m+2, G), fumarate (m+2, H), and malate (m+2, I) in WT and MLL3/4 DKO ESCs labeled with [^13^C_6_]-glucose. n=3 biological replicates per sample. Data are presented as mean ± S.D. p-values were calculated using Student’s t-test. (J) Schematic of the TCA cycle illustrating the fate of ^13^C carbons upon the incubation with [^13^C_5_]-glutamine. (K-P) Bar plots showing levels of glutamate (m+5, K), α-KG (m+5, L), succinate (m+4, M), fumarate (m+4, N), malate (m+4, O), and citrate/isocitrate (m+4, P) in WT and MLL3/4 DKO ESCs labeled with [^13^C_5_]-glutamine. n=3 biological replicates per sample. Data are presented as mean ± S.D. p-values were calculated using Student’s t-test.(Q-T) Western blots showing the levels of OGDH (Q), DLST (R), DLD (S), and lipoylated proteins (T) in WT and MLL3/4 DKO ESCs. 10 and 20 μg of cell lysates were loaded for each sample. Experiments were repeated three times independently and representative results are shown. Protein levels of OGDH (in Q), DLST (in R), DLD (in S), and lipoylated DLST (in T) were quantified by normalizing with Tubulin or HSP90 signals of the corresponding lane.

To further test this model, we traced glutamine carbon entry into the TCA cycle using [^13^C_5_]-glutamine (Fig. 2J). Although MLL3/4 loss did not significantly affect labeling of glutamate (m+5) (Fig. 2K and Fig. S2F), that of α-KG (m+5) was significantly reduced in MLL3/4 DKO ESCs (Fig. 2L and Fig. S2G). This reduction may reflect impaired conversion of glutamate to α-KG, although the expression levels of *Glud1*, which encodes glutamate dehydrogenase (GDH), the enzyme catalyzing the conversion of glutamate to α-KG, were comparable between WT and MLL3/4 DKO ESCs (Fig. S2L). Consistent with results from [^13^C_6_]-glucose labeling, labeling of succinate (m+4), fumarate (m+4), and malate (m+4) from [^13^C_5_]-glutamine was significantly reduced in MLL3/4 DKO ESCs, while that of citrate/isocitrate (m+4) was largely unchanged (Fig. 2M-P and Fig. S2H-K). Furthermore, expression levels of multiple genes encoding enzymes involved in α-KG and downstream TCA cycle metabolism were comparable between MLL3/4 DKO and WT ESCs (Fig. S2L). In addition, protein levels of OGDH, SUCLA2, SUCLG1, SUCLG2, and SDHA remained largely unperturbed upon MLL3/4 loss (Fig. 2Q and Fig. S2M-N), suggesting that MLL3/4 regulate TCA cycle metabolites via mechanisms other than transcriptional control of metabolic enzymes.

To further elucidate the molecular mechanisms by which MLL3/4 regulate the TCA cycle, we examined the protein levels of DLST and dihydrolipoamide dehydrogenase (DLD), E2 and E3 components of the OGDHc. MLL3/4 loss had little impact on their levels (Fig. 2R-S), suggesting that MLL3/4 are unlikely to regulate mitochondrial respiration via modulating the abundance of OGDHc components. Lipoylated DLST is essential for OGDHc activity and for the conversion of α-KG to succinyl-CoA (*23-25*). To understand if MLL3/4 loss impairs the activity of DLST, we examined the levels of lipoylated proteins in WT and MLL3/4 DKO ESCs using a specific anti-lipoate antibody (*26*). MLL3/4 loss led to a reproducible decrease of the lipoylated protein species with lower molecular weight, attributed to lipoylated DLST based on its size (Fig. 2T). The decrease in lipoylation was specific to DLST as levels of other abundantly lipoylated proteins in the mitochondria, such as dihydrolipoamide acetyltransferase (DLAT) in the pyruvate dehydrogenase complex and dihydrolipoamide branched-chain transacylase (DBT) in the branched-chain alpha-keto acid dehydrogenase complex (*24, 25*), were comparable in MLL3/4 DKO and WT ESCs (Fig. 2T). Taken together, these results suggest that MLL3/4 loss impairs mitochondrial respiration and disrupts metabolic flux through reducing DLST lipoylation in naive PSCs.

### Combined overexpression of HK2 and OGDH rescues metabolic defects caused by MLL3/4 loss

To determine if MLL3/4 regulate glycolysis and mitochondrial respiration in ESCs through activating HK2, we overexpressed 3×HA-tagged HK2 in MLL3/4 DKO ESCs with a doxycycline (DOX)-inducible system (*27, 28*) (Fig. S3A). HK2 overexpression increased basal glycolysis and glycolytic capacity in MLL3/4 DKO cells (Fig. S3B-D); however, OCR during basal respiration was reduced upon HK2 expression (Fig. S3E-G), suggesting that HK2 overexpression renders MLL3/4 DKO ESCs more dependent on glycolysis for energy production.

Since HK2 overexpression led to an expected increase in ECAR but a reduction in OCR, we sought to restore both glycolysis and mitochondrial respiration in MLL3/4 DKO ESCs to study the functional relationship of MLL3/4, metabolic state, and the differentiation capability of PSCs. We hypothesized that the decrease of ^13^C incorporation in succinate, fumarate, and malate in both [^13^C_6_]-glucose and [^13^C_5_]-glutamine labelled MLL3/4-null ESCs results from the defective OGDHc. Although OGDH levels were comparable between MLL3/4 DKO and WT ESCs, OGDH overexpression could potentially compensate for the functionally impaired OGDHc and reduced TCA cycle metabolites in MLL3/4-null ESCs. We therefore overexpressed 3×HA-tagged HK2 and 3×FLAG-tagged OGDH together in MLL3/4 DKO ESCs using DOX-inducible expression vectors carrying distinct drug resistance genes (Fig. 3A). Increased levels of HK2 and OGDH led to elevated rates of basal glycolysis, glycolytic capacity, and both basal and maximal respiration (Fig. 3B-G). Furthermore, stable isotope tracing by labeling MLL3/4 DKO and HK2/OGDH double-overexpressing (DOE) MLL3/4 DKO cells with [^13^C_6_]-glucose indicated that labeling of metabolites reduced by MLL3/4 loss such as F6P/G1P/G6P (m+6), lactate (m+3), succinate (m+2), and fumarate (m+2) was partially restored by HK2/OGDH DOE (Fig. 3H-M, Fig. S4A-B, and Fig. S4E-G). As expected, we did not observe a significant difference in the labeling of citrate/isocitrate (m+2) or α-KG (m+2) (Fig. 3J and Fig. S4C-E). Notably, malate (m+2) labeling was not significantly increased upon HK2/OGDH DOE in MLL3/4 DKO cells (Fig. 3M and Fig. S4H), suggesting that there are additional regulatory layers on the levels of malate besides HK2 and OGDHc. Taken together, these results suggest that MLL3/4 regulate glycolysis and mitochondrial respiration through activating *Hk2* expression and maintaining the normal function of the OGDHc.

**Figure 3.**
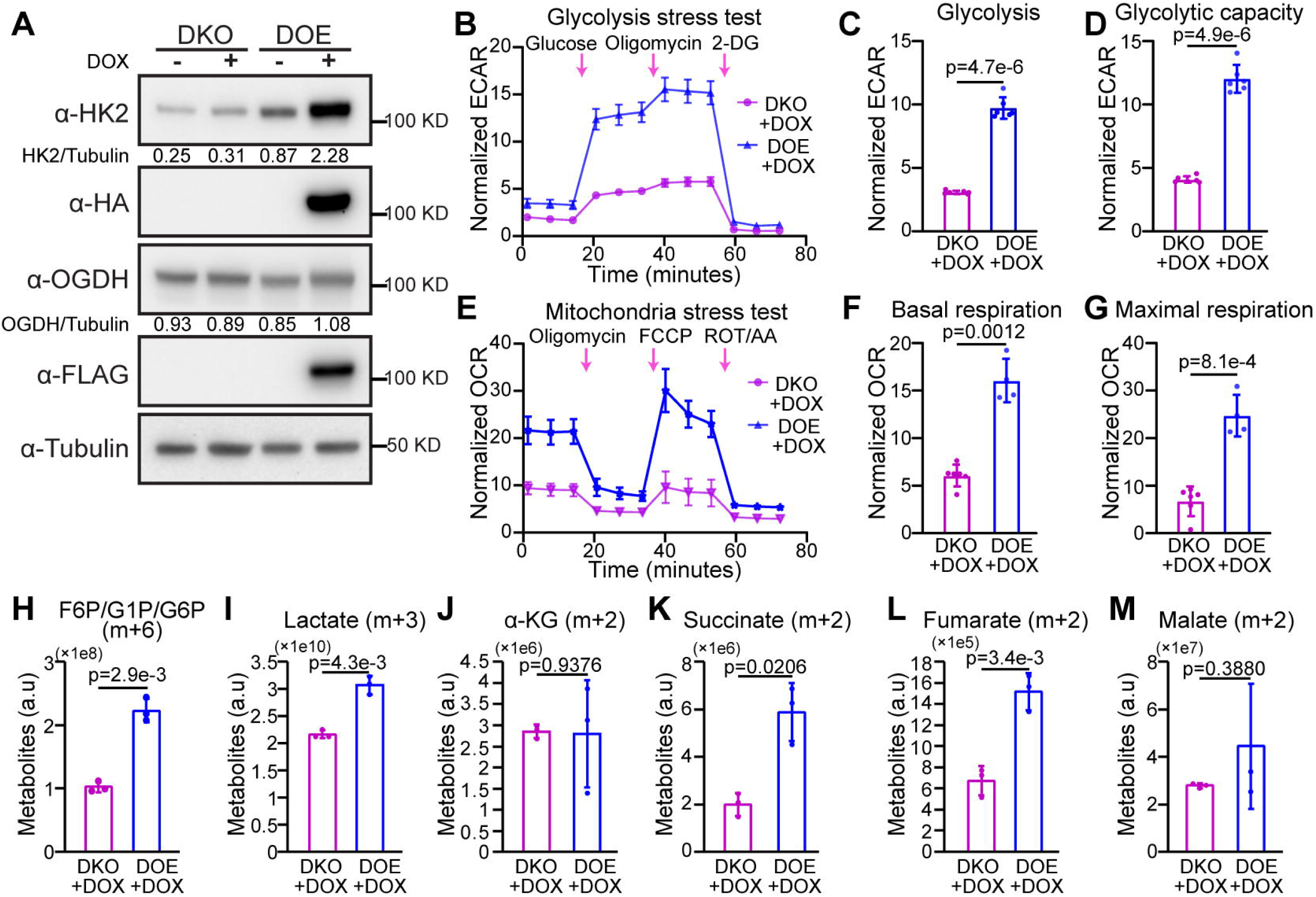
Combined overexpression of HK2 and OGDH rescues metabolic defects of MLL3/4 DKO ESCs. (A) Western blots showing levels of HK2, 3×HA-tagged HK2, OGDH, and 3×FLAG-tagged OGDH in MLL3/4 DKO ESCs and MLL3/4 DKO ESCs overexpressing HK2/OGDH treated with or without DOX. 10 and 20 μg of cell lysates were loaded for each sample. Experiments were repeated three times independently and representative results are shown. Protein levels in each lane of the anti-HK2 and anti-OGDH blots were quantified by normalizing with Tubulin signals of the corresponding lane. (B) Representative ECAR traces of MLL3/4 DKO ESCs treated with DOX (DKO+DOX) and MLL3/4 DKO ESCs overexpressing HK2/OGDH (DOE+DOX) measured by the Seahorse XF glycolysis stress test assay. Data were normalized by the number of cells at the time of the assay and are represented as mean ± S.D. Each data point had six replicates, and the experiment was repeated three times independently. (C-D) Bar plots of ECAR showing the level of glycolysis (C) and glycolytic capacity (D) in MLL3/4 DKO ESCs treated with DOX (DKO+DOX) and MLL3/4 DKO ESCs overexpressing HK2/OGDH (DOE+DOX). Data are presented as mean ± S.D. p-values were calculated using Student’s t-test. (E) Representative OCR traces of MLL3/4 DKO ESCs treated with DOX (DKO+DOX) and MLL3/4 DKO ESCs overexpressing HK2/OGDH (DOE+DOX) measured by the Seahorse XF mitochondrial stress test assay. Data were normalized by the number of cells at the time of the assay and are represented as mean ± S.D. Each data point had six replicates, and the experiment was repeated three times independently. (F-G) Bar plots showing the levels of OCR during basal (F) and maximal (G) respiration in MLL3/4 DKO ESCs treated with DOX (DKO+DOX) and MLL3/4 DKO ESCs overexpressing HK2/OGDH (DOE+DOX). Data are presented as mean ± S.D. p-values were calculated using Student’s t-test. (H-M) Bar plots showing levels of F6P/G1P/G6P (m+6, H), lactate (m+3, I), α-KG (m+2, J), succinate (m+2, K), fumarate (m+2, L), and malate (m+2, M) in MLL3/4 DKO ESCs treated with DOX (DKO+DOX) and MLL3/4 DKO ESCs overexpressing HK2/OGDH (DOE+DOX) labeled with [^13^C_6_]-glucose. n=3 biological replicates per sample. Data are presented as mean ± S.D. p-values were calculated using Student’s t-test.

### Combined overexpression of HK2 and OGDH rescues differentiation defects caused by MLL3/4 loss

To assess the impact of MLL3/4 loss on cell fate transition, we generated embryoid bodies (EBs) from WT and MLL3/4 DKO ESCs. As expected, MLL3/4 loss led to a decrease in the size of day 6 EBs (Fig. 4A-B), indicating that MLL3/4 are required for PSC differentiation. RNA-seq analysis showed that MLL3/4 loss caused the significant deregulation of 4,547 genes in day 6 EBs (Fig. 4C). Among genes significantly downregulated in MLL3/4 DKO EBs, 52.3% (1,192 of 2,280) were significantly upregulated during WT EB differentiation (Fig. 4D), suggesting that these genes are dependent on MLL3/4 to be activated during differentiation. We thus defined them as MLL3/4-dependent differentiation genes. Gene ontology (GO) analysis showed that the top 300 upregulated genes of the 1,192 MLL3/4-dependent differentiation genes were enriched with embryogenesis-related genes and known MLL4 targets essential for actin cytoskeleton organization (*7*) (Fig. 4E), further indicating that MLL3/4 loss impairs PSC differentiation. Indeed, pluripotency genes were significantly upregulated in MLL3/4 DKO EBs and marker genes of meso- and endo-derm lineages were downregulated in EBs by MLL3/4 loss (Fig. S5A-B). These data demonstrate that MLL3/4 are required for PSC differentiation and related transcriptional reconfiguration.

**Figure 4.**
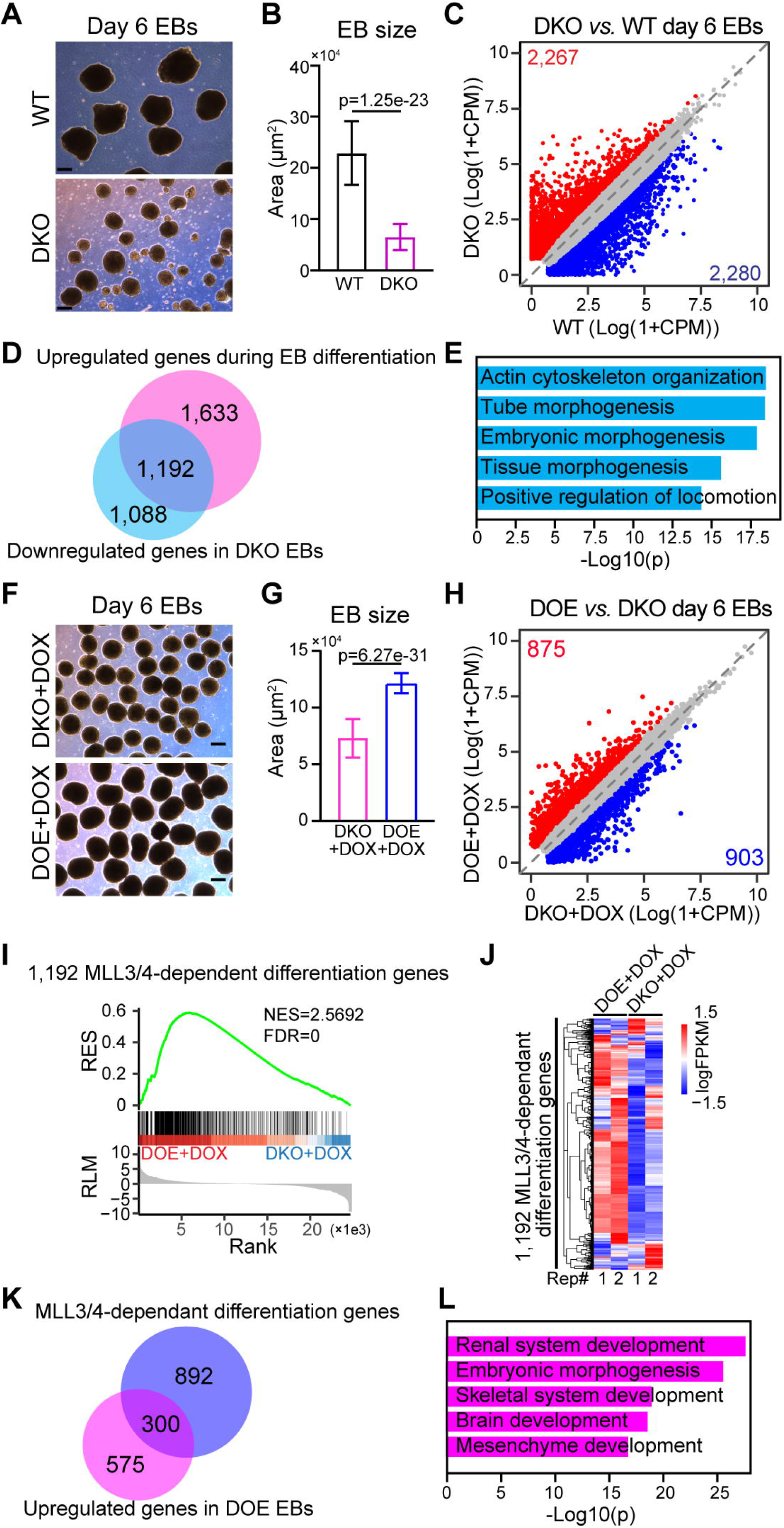
Combined overexpression of HK2 and OGDH rescues differentiation defects caused by MLL3/4 loss. (A) Representative images of day 6 WT and MLL3/4 DKO EBs. Scale bar: 250μm. (B) Quantification of EB sizes in (A). n=50 for each genotype. Experiments were repeated three times independently and representative results are shown. (C) Correlation plots of RNA-seq data between day 6 MLL3/4 DKO and WT EBs. Data were derived from two biological replicates. Statistical significance was determined by two-sided Wald test and p-values were corrected for multiple testing using the Benjamini–Hochberg method. Significantly up- and downregulated genes (adjusted p<0.01, ΙlogFCΙ>1) were labeled in red and blue with numbers of genes noted, respectively. (D) Venn diagram identifying 1,192 MLL3/4-dependent differentiation genes. (E) GO analysis of top 300 MLL3/4-dependent differentiation genes. (F) Representative images of day 6 MLL3/4 DKO EBs treated with DOX (DKO+DOX) and MLL3/4 DKO EBs overexpressing HK2/OGDH (DOE+DOX). Scale bar: 250μm. (G) Quantitation of EB sizes in (F). n=50 for each genotype. Experiments were repeated three times independently and representative results are shown. (H) Correlation plots of RNA-seq data between MLL3/4 DKO EBs treated with DOX (DKO+DOX) and MLL3/4 DKO EBs overexpressing HK2/OGDH (DOE+DOX). (I) GSEA analysis of MLL3/4-dependent differentiation genes (1,192 in D) comparing MLL3/4 DKO EBs treated with DOX (DKO+DOX) and MLL3/4 DKO EBs overexpressing HK2/OGDH (DOE+DOX). RES: running enrichment score; RLM: ranked list metric; NES: normalized enrichment score; FDR: false discovery rate. (J) Hierarchical clustering analysis of the expression levels of 1,192 MLL3/4-dependent differentiation genes in MLL3/4 DKO EBs treated with DOX (DKO+DOX) and MLL3/4 DKO EBs overexpressing HK2/OGDH (DOE+DOX). FPKM: Fragments per kilobase of transcript per million mapped reads. (K) Venn diagram showing the overlap of MLL3/4-dependent differentiation genes and significantly upregulated genes in MLL3/4 DKO EBs overexpressing HK2/OGDH (red in H). (L) GO analysis of 300 DOE-rescued differentiation genes (overlapping genes in K).

To examine if the partial restoration of glycolysis and mitochondrial respiration had any impact on the differentiation capacity of ESCs lacking MLL3/4, we treated MLL3/4 DKO and HK2/OGDH DOE MLL3/4 DKO cells with DOX and differentiated them toward EBs. EB differentiation revealed that HK2/OGDH DOE led to a significant increase in the size of day 6 EBs (Fig. 4F-G), suggesting that restoring glycolysis and mitochondrial respiration rescues differentiation defects caused by MLL3/4 loss. Moreover, RNA-seq analysis identified 1,778 significantly deregulated genes by HK2/OGDH DOE compared with MLL3/4 DKO ESCs treated with DOX (Fig. 4H). Gene set enrichment analysis (GSEA) and hierarchical clustering analysis of MLL3/4-dependent differentiation genes (Fig. 4D) revealed a significant upregulation of them upon HK2/OGDH DOE (Fig. 4I-J). Indeed, 34.3% of genes upregulated by HK2/OGDH DOE (300 of 875) were dependent on MLL3/4 to be activated during differentiation (Fig. 4K), suggesting that the impaired metabolic state contributes to differentiation defects caused by MLL3/4 loss. GO analysis of the 300 significantly rescued genes showed enrichment of genes related to embryo morphogenesis and mesodermal development such as the development of renal and skeletal systems (Fig. 4L). Consistently, HK2/OGDH DOE led to the significant downregulation of pluripotency genes and the significant upregulation of differentiation marker genes (Fig. S5C-D). Taken together, these results suggest that defects in glycolysis and mitochondrial respiration contribute to differentiation failures of naive PSCs lacking MLL3/4.

## Discussion

Epigenetic regulators, including histone modifiers, play central roles in regulating cell fate transition during PSC differentiation, somatic cell reprogramming, and normal development (*29-35*); however, the underlying mechanisms remain incompletely understood, in part because many epigenetic regulators function independently from their catalytic activity (*6, 7, 28, 36-43*). Besides being essential for cellular differentiation and mammalian embryogenesis (*11, 44*), loss-of-function mutations in MLL3 and MLL4 drive congenital disorders such as Kleefstra syndrome 2 and Kabuki syndrome, as well as multiple types of cancer (*1, 4, 13, 45*). Recent advances reveal a novel link between MLL3/4 and cellular metabolism; however, whether and how the roles of MLL3/4 in regulating cell fate and metabolic pathways intersect are unclear. Here, we unveil a previously unrecognized function of MLL3/4 in regulating glycolysis and mitochondrial respiration in murine ESCs. Our findings suggest that MLL3/4 govern cell fate transition during the differentiation of naive PSCs through regulating the expression levels of *Hk2* and the function of OGDHc, two rate-limiting nodes in glycolysis and the TCA cycle, respectively. Importantly, restoring defective glycolysis and mitochondrial respiration may be an effective strategy to treat diseases driven by MLL3/MLL4 loss-of-function.

We find that MLL3/4 loss causes decreased glycolysis and mitochondrial respiration in ESCs. Metabolic bivalency is a defining feature of naive PSCs such as murine ESCs (*22*). The decrease of glycolysis in MLL3/4-null cells may hinder the cells from exiting naive pluripotency, as formative PSCs and primed PSCs are highly glycolytic (*17*). On the other hand, inhibiting glycolysis promotes differentiation as somatic cells have much elevated levels of oxidative phosphorylation compared with PSCs (*18-20*). Therefore, the dampened mitochondrial respiration caused by MLL3/4 loss may impair the differentiation of PSCs into germ layers and/or specific lineages. Our data provide a mechanistic explanation for the failure of MLL4-null ESCs to exit naive pluripotency and for the severe gastrulation defects observed in MLL3/4-null embryos (*7, 10, 46*).

Our data support that MLL3/4 facilitate glycolysis in naive ESCs through maintaining the expression of *Hk2*, which encodes the rate-limiting enzyme hexokinase 2 in glycolysis. It is possible that MLL3/4 regulate *Hk2* expression through enhancers as we identified multiple putative *cis*-regulatory regions that may be directly regulated by MLL3/4. Understanding whether and how MLL3/4 function through these putative enhancers to control the expression levels of *Hk2* is an interesting topic. Shadow enhancers have been reported in different organisms including *Drosophila*, mouse, and human (*47-49*). Generating individual and compound enhancer mutations will elucidate how the putative enhancers regulate *Hk2*. Combination of MLL3/4 loss-of-function and enhancer mutations will further reveal the functional relationship between MLL3/4 and putative enhancers in regulating *Hk2*. These future studies are essential for understanding the mechanisms by which MLL3/4 regulate glycolysis in naive PSCs.

Interestingly, overexpressing HK2 alone only partially restores glycolysis in ESCs lacking MLL3/4, suggesting that MLL3/4 support glycolysis in naive PSCs through additional mechanisms. Determining whether other critical regulators of glycolysis such as phosphofructokinase, pyruvate kinase, and lactate dehydrogenase play a role in dampening glycolysis in the context of MLL3/4 loss-of-function is an important future study. Genome-wide and targeted genetic screens may also identify additional factors that rescue glycolysis in MLL3/4-null ESCs. It is noteworthy that HK2 overexpression leads to a decrease of OCR in MLL3/4 DKO ESCs, suggesting that these cells are more dependent on glycolysis as an energy source in comparison to oxidative phosphorylation under high levels of HK2. Since steady-state acetyl-CoA abundance was reduced in MLL3/4 DKO ESCs (*14*) and our stable isotope tracing assays were not able to detect labeled acetyl-CoA, it is interesting to examine if and how MLL3/4 loss affects the mitochondrial transport of pyruvate in ESCs in the future. Defining the role of MLL3/4 in other pathways leading to the synthesis of acetyl-CoA such as fatty acid metabolism may also facilitate understanding how MLL3/4 regulate the conjunction of glycolysis and the TCA cycle.

The reduction in ^13^C incorporation into succinate, fumarate, and malate in both [^13^C_6_]-glucose and [^13^C_5_]-glutamine labeling of MLL3/4 DKO ESCs suggests a functionally defective OGDHc caused by MLL3/4 loss. Indeed, the levels of lipoylated DLST were decreased in MLL3/4-null cells, suggesting a reduced activity of the OGDHc. It is likely that the decrease of lipoylation is specific to DLST as MLL3/4 loss has little impact on the levels of lipoylated DLAT. It was previously shown that the α/β hydrolase ABHD11 binds OGDHc and maintains the activity of lipoylated DLST by preventing the formation of lipoyl adducts (*26*). It will be interesting to investigate the functional relationship of MLL3/4, ABHD11, and OGDHc in the future to understand how MLL3/4 regulate the activity of OGDHc. It is also noteworthy that ^13^C incorporation into α-KG (m+5) is impaired in MLL3/4 DKO ESCs labelled with [^13^C_5_]-glutamine, although α-KG (m+2) levels through [^13^C_5_]-glucose labeling and steady-state levels of α-KG were largely unchanged by MLL3/4 loss (*14*). Besides glutamate dehydrogenase, glutamate pyruvate transaminase and glutamate oxaloacetate transaminase also can convert glutamate to α-KG. MLL3/4 may also regulate the transport of cytosolic glutamate into mitochondria. Future studies on the role of MLL3/4 in regulating glutamate transport and the conversion of glutamate to α-KG would be important to understand the crosstalk between MLL3/4 and glutaminolysis.

α-KG is an essential co-substrate of Fe(II)/α-KG-dependent dioxygenases including Jumonji C (JmjC) domain-containing histone demethylases and the ten-eleven translocation (TET) family of DNA hydroxylases (*50, 51*), playing a role in regulating the levels of methylation on different lysine residues of histones and 5-methyl Cytosine (5mC) on DNA in mammalian cells. Due to their structural similarity to α-KG, 2-hydroxyglutarate, succinate, and fumarate inhibit α-KG-dependent dioxygenases by acting as competitive inhibitors of α-KG (*52, 53*). Our results that MLL3/4 loss in ESCs leads to decreased levels of succinate and fumarate, through steady-state metabolomics (*14*) and stable isotope tracing with [^13^C_6_]-glucose and [^13^C_5_]-glutamine, suggest a possibility that activities of multiple histone demethylases and TETs are increased in cells with MLL3/4 loss-of-function. Such a possibility provides a mechanistic explanation for the DNA hypomethylation observed in MLL4-depleted B cells in mouse and MLL4 mutant prostate cancers in human (*54*). Understanding the impact of the potential alteration in levels of histone methylation and DNA methylation on the differentiation capability of MLL3/4 mutant PSCs is very interesting and may provide insight into understanding the molecular mechanisms underlying the functional relationship of metabolic state, epigenetic machineries, and cell fate transition.

In summary, our study reveals a previously unappreciated crosstalk between epigenetic modifiers exemplified by MLL3/4 and glycolysis and mitochondrial respiration in regulating cell fate. Our findings on MLL3/4 regulating the levels of HK2 and lipoylated DLST could pave the way for designing therapeutic strategies to target human diseases driven by loss-of-function mutations of MLL3/4.

## Methods

### Antibodies

The following primary antibodies were used in this study: anti-HK2 (Cell Signaling Technology 2867), anti-HK1 (Cell Signaling Technology 2024), anti-OGDH (Proteintech 17110-1-AP), anti-SDHA (Abcam 14715), anti-SUCLA2 (Proteintech 12627-1-AP), anti-SUCLG1 (Proteintech 14923-1-AP), anti-SUCLG2 (Proteintech NBP1-32521), anti-FLAG (MiliporeSigma F3165), anti-HA (MiliporeSigma H3663), anti-DLST (Cell Signaling Technology 5556), anti-DLD (Santa Cruz 365977), anti-Lipoic acid (Calbiochem 437695), anti-Tubulin (Developmental Studies Hybridoma Bank E7), and anti-HSP90 (Santa Cruz 13119).

### ESC Culture and EB differentiation

ESCs were cultured in serum-free medium supplemented with 2i/LIF as previously described (*55*). For the generation of HK2 overexpressing and HK2/OGDH DOE MLL3/4 DKO ESCs, cells were transfected with DOX-inducible PiggyBac vectors (*27*) as previously described (*28*) using an Amaxa Nucleofector (Lonza). EB differentiation was performed using the hanging-drop method as previously described (*28*). For DOX-induced overexpression, ESCs were treated with 100 ng/ml doxycycline for 48 hours prior to downstream assays. For differentiation experiments, doxycycline (100 ng/ml) were added 24 hours prior to EB formation and maintained throughout the 6-day differentiation period.

### Seahorse extracellular flux assays

ESCs were seeded in Seahorse XF microplates (Agilent) and cultured overnight with 2i/LIF serum-free medium. Cells were then incubated in Seahorse XF assay medium (Agilent) at 37°C in a non-CO_2_ incubator for 1 hour. For the Glycolysis Stress Test (Agilent 103020-100), glucose (10 mM), oligomycin (2 µM), and 2-DG (50 mM) were sequentially injected. For the Mitochondrial Stress Test (Agilent 103015-100), oligomycin (1.5 µM), FCCP (0.5 µM), and Rot/AA (0.5 µM) were sequentially injected. Cell Titer Glo assay (Promega) was performed in parallel on replica plates to determine the cell number of each well. Individual raw ECAR and OCR values recorded by the Seahorse XF Analyzer were normalized against the average Cell Titer Glo values of control wells to derive normalized ECAR and normalized OCR values. Wave software (Agilent) was used to extract individual ECAR and OCR measurements, which were subsequently used to generate the bar plots.

### Stable isotope tracing and metabolomics

Stable isotope labeling and metabolite extraction were performed as previously described (*14*). ESCs were incubated with media containing 25 mM [^13^C_6_]-glucose or 4 mM [^13^C_5_]-glutamine for 2 hours before methanol extraction. Metabolomic analyses were performed at the Northwestern Metabolomics Core Facility.

### Glucose uptake assay

Glucose update was measured using the Glucose Uptake-Glo Assay Kit (Promega J1342) according to the manufacturer’s instructions. In brief, 2×10^4^ cells were seeded on each well of a 96-well flat bottom plate with opaque walls and clear bottoms (Corning 7200587). After 16 hours of culture, cells were incubated in glucose-free medium for 1 hour at 37°C in a CO_2_ incubator. Cells were then treated with 1 mM 2-DG for 10 minutes at room temperature, followed by the treatment of stop buffer, neutralization buffer, and detection reagent following the instructions of the manufacturer. After 2.5 hours of dark-conditioned incubation, the luminescence was measured using a GloMax luminometer (Promega). The luminescence data were normalized by cell numbers measured with the Cell Titer Glo assay (Promega) on replica plates in parallel.

### RNA-seq and analysis

Total RNA was extracted using TRIzol reagent (Thermo Fisher) following the manufacturer’s instructions. Each cell line was analyzed in at least two biological replicates. Extracted RNA were treated with DNase I (Sigma) and purified with the RNeasy Plus Mini Kit (Qiagen). Ribosomal RNA was removed from the purified RNA with the NEBNext rRNA Depletion Kit (New England BioLabs). RNA libraries were then prepared with the NEBNext Ultra II Directional RNA Library Prep Kit (New England BioLabs). The pooled libraries were sequenced with 150 bp paired-end reads on the NovaSeq platfrom (Illumina).

Raw reads were processed with Trimmomatic v0.39(*56*) to remove adaptors and low-quality reads with the parameter “-q 30”. The cleaned reads were aligned to the mm9 genome assembly using STAR v2.5.1 with default parameter (*57*). Reads were normalized to total read counts per million (cpm) and visualized as bigwig-formatted coverage tracks using deepTools v3.5.6 (*58*). Gene expression was quantified using HTseq V2.0.2 (*59*), and differential gene expression analysis was performed with EdgeR v4.0.16 (*60*). Significantly differentially expressed genes were identified with Benjamini-Hochberg-adjusted p value cutoff < 0.01 and a log fold change threshold greater than |1|. hierarchical clustering heatmaps were generated using the ggpubr v0.6.0 (https://rpkgs.datanovia.com/ggpubr/). GSEA plots were generated with clusterProfiler v4.10.1 (*61*). Gene ontology (GO) analysis was performed using Metascape v3.5.20260201 (*62*).

## Acknowledgements

We thank the Cao lab members for their helpful suggestions and discussions. We thank Dr. Peng Gao and the Northwestern Metabolomics Core Facility for performing the metabolomics study. We thank Emmalee W. Cooke for technical support. We thank Yalu Zhou for bioinformatic analysis support. This study was supported in part by grants R35GM150668, R00HD094906, and P30CA043703 from the National Institutes of Health to K.C.

## Author contributions

K.C. conceived the study. S.M.N., I.B., and K.C. designed the research. S.M.N., Y.J., and M.Y. performed experiments. S.M.N., Y.J., M.Y., and K.C. analyzed the data. S.M.N. and K.C. wrote the manuscript. C.A.L. and I.B. critically read the manuscript.

## Figure legends

**Figure S1. The impact of MLL3/4 loss on glycolytic metabolites, glycolytic gene expression, and putative enhancers at the *Hk2* locus.**

(A-B) Stable isotope tracing of WT and MLL3/4 DKO ESCs labeled with [^13^C_6_]-glucose. The levels of isotopologues (m+0 to m+6) of F6P/G6P/G1P (A) and lactate (B) were shown as stacked bar plots.

(C) Relative expression levels of genes coding key glycolytic enzymes involved in the production and utilization of G6P, F6P, and lactate. CPM (counts per million reads) from RNA-seq data (*14*) of MLL3/4 DKO ESCs were normalized to those of WT ESCs. p-values were determined by two-sided Wald test and corrected using the Benjamini–Hochberg method.

(D) Genome browser view of MLL4 ChIP-seq signals in WT ESCs, and that of H3K27ac and H3K4me3 ChIP-seq signals in WT and MLL3/4 DKO ESCs at the *Hk2* locus. Putative enhancers bound by MLL4 around *Hk2* gene were highlighted.

**Figure S2. The impact of MLL3/4 loss on TCA cycle metabolites and the expression of TCA cycle-related genes.**

(A-E) Stable isotope tracing of WT and MLL3/4 DKO ESCs labeled with [^13^C_6_]-glucose. The levels of isotopologues (m+0 to m+6) of citrate/isocitrate (A), α-KG (B), succinate (C), fumarate (D), and malate (E) were shown as stacked bar plots.

(F-K) Stable isotope tracing of WT and MLL3/4 DKO ESCs labeled with [^13^C_5_]-glutamine. The levels of isotopologues (m+0 to m+6) of glutamate (F), α-KG (G), succinate (H), fumarate (I), malate (J), and citrate/isocitrate (K) were shown as stacked bar plots.

(L) Relative expression levels of genes coding TCA-cycle enzymes involved in the production and utilization of α-KG, succinate, fumarate, and malate. CPM (counts per million reads) from RNA-seq data (*14*) of MLL3/4 DKO ESCs were normalized to those of WT ESCs. p-values were determined by two-sided Wald test and corrected using the Benjamini–Hochberg method.

(M-N) Western blots showing levels of Succinyl-CoA Synthetase subunits (M), SDHA (N, top blot), and FH (N, middle blot). 10 and 20 μg of cell lysates were loaded for each sample. Experiments were repeated three times independently and representative results are shown. Protein levels in each lane were quantified by normalizing with HSP90 signals of the corresponding lane.

**Figure S3. HK2 overexpression leads to an increase in glycolysis but a decrease in mitochondrial respiration in MLL3/4 DKO ESCs.**

(A) Western blots showing the induction of 3×HA-tagged HK2 in HK2-overexpressing MLL3/4 DKO ESCs by DOX. 10 and 20 μg of cell lysates were loaded for each sample. Experiments were repeated three times independently and representative results are shown. Protein levels of HK2 in each lane were quantified by normalizing with Tubulin signals of the corresponding lane.

(B) Representative ECAR traces of MLL3/4 DKO ESCs treated with DOX (DKO+DOX) and MLL3/4 DKO ESCs overexpressing HK2 (3×HA-HK2+DOX) measured by the Seahorse XF glycolysis stress test assay. Data were normalized by the number of cells at the time of the assay and are represented as mean ± S.D. Each data point had six replicates, and the experiment was repeated three times independently.

(C-D) Bar plots of ECAR showing the levels of glycolysis (C) and glycolytic capacity (D) in MLL3/4 DKO ESCs treated with DOX (DKO+DOX) and MLL3/4 DKO ESCs overexpressing HK2 (3×HA-HK2+DOX). Data are presented as mean ± S.D. p-values were calculated using Student’s t-test.

(E) Representative OCR traces of MLL3/4 DKO ESCs treated with DOX (DKO+DOX) and MLL3/4 DKO ESCs overexpressing HK2 (3×HA-HK2+DOX) measured by the Seahorse XF mitochondrial stress test assay. Data were normalized by the number of cells at the time of the assay and are represented as mean ± S.D. Each data point had six replicates, and the experiment was repeated three times independently.

(F-G) Bar plots showing the OCR levels in MLL3/4 DKO ESCs treated with DOX (DKO+DOX) and MLL3/4 DKO ESCs overexpressing HK2 (3×HA-HK2+DOX) during basal (F) and maximal

(G) respiration. Data are presented as mean ± S.D. p-values were calculated using Student’s t-test.

**Figure S4. Stable isotope tracing of MLL3/4 DKO ESCs treated with DOX (DKO+DOX) and MLL3/4 DKO ESCs overexpressing HK2/OGDH (DOE+DOX) labeled with [^13^C_6_]-glucose.**

The levels of isotopologues of F6P/G1P/G6P (A, m+0 to m+6), lactate (B m+0 to m+3), citrate/isocitrate (m+2 in C and m+0 to m+6 in D), α-KG (E, m+0 to m+5), succinate (F, m+0 to m+4), fumarate (G, m+0 to m+4), and malate (H, m+0 to m+4) were shown.

**Figure S5. The impact of MLL3/4 loss and HK2/OGDH overexpression on the expression levels of marker genes of pluripotency and differentiation in EBs.**

(A-B) Relative expression levels of pluripotency genes (A) and differentiation markers (B) in MLL3/4 DKO EBs compared with WT EBs. Log2 fold change of RNA-seq signals (CPM) were shown. Data are presented as mean values ± SD. n = 2 biologically independent experiments. All p-values were smaller than 0.01 compared with WT EBs.

(C-D) Relative expression levels of pluripotency genes (C) and differentiation markers (D) in MLL3/4 DKO EBs overexpressing HK2/OGDH (DOE+DOX) compared with MLL3/4 DKO EBs treated with DOX (DKO+DOX). Log2 fold change of RNA-seq signals (CPM) were shown. Data are presented as mean values ± SD. n = 2 biologically independent experiments. All p-values were smaller than 0.01 compared with DKO+DOX EBs.

