## Supplementary figures and images for "MLL3/4 methyltransferases regulate the differentiation of pluripotent stem cells through coordinating glycolysis and mitochondrial respiration"

### Figure S1

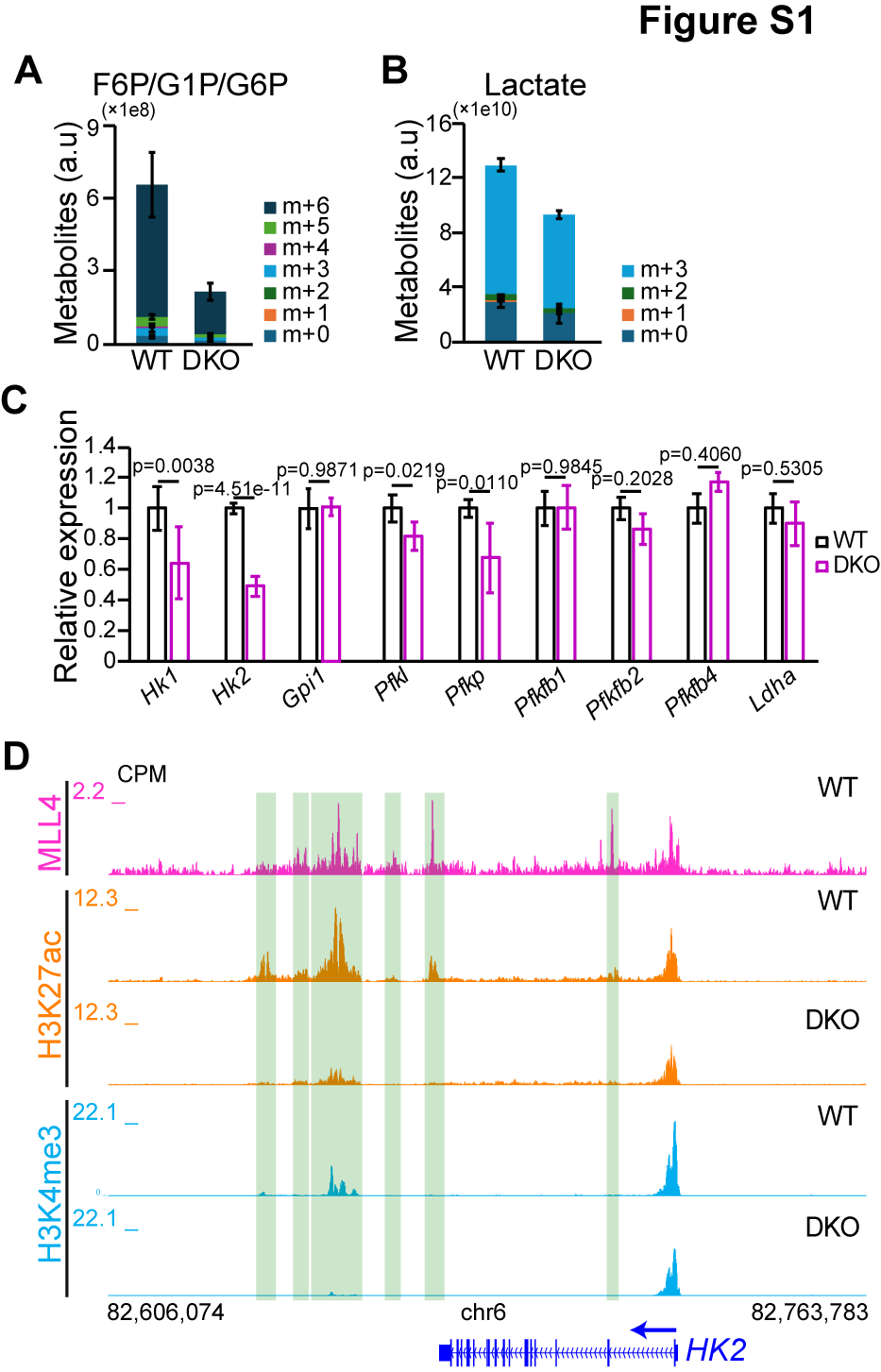

### Figure S2

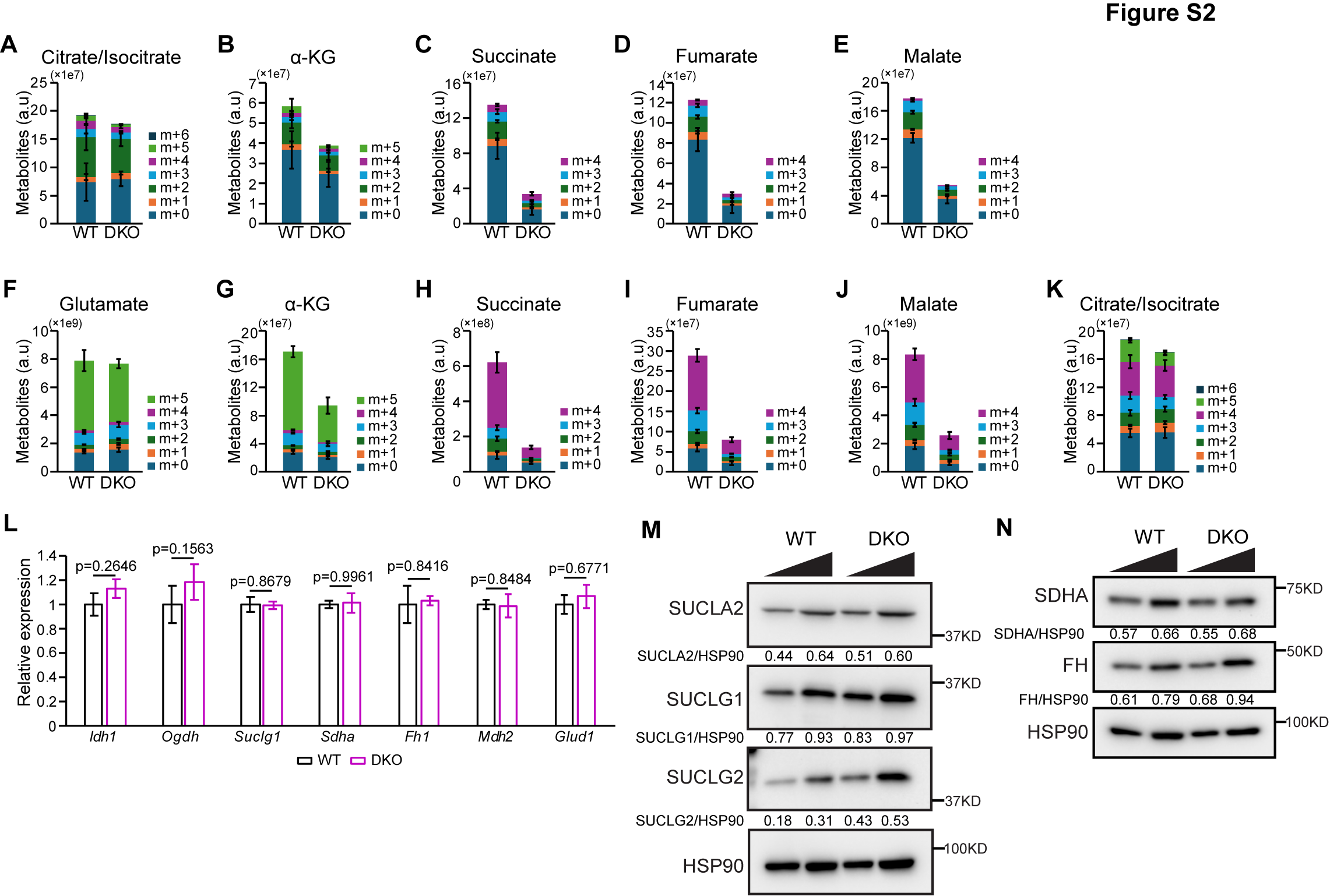

### Figure S3

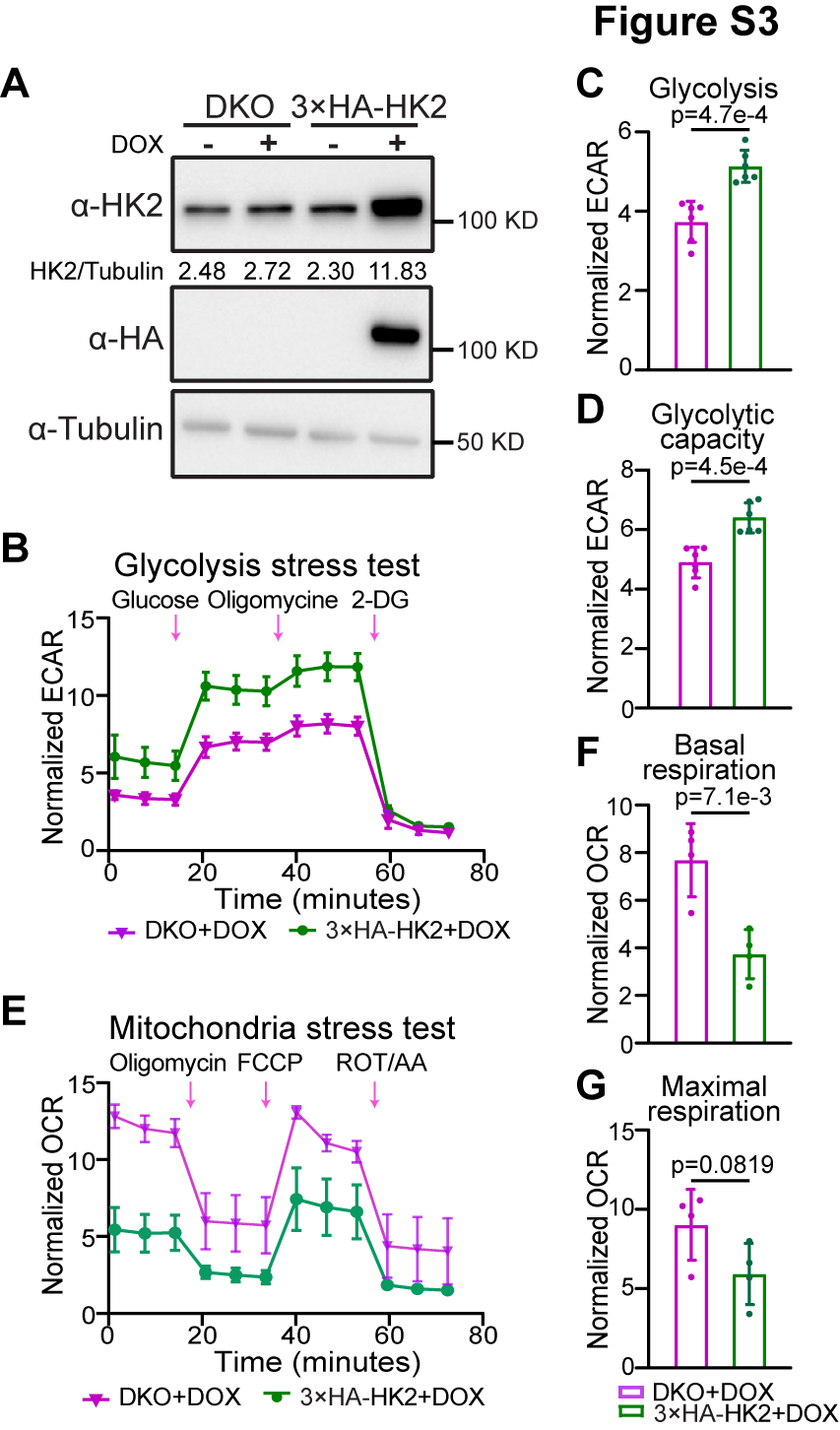

### Figure S4

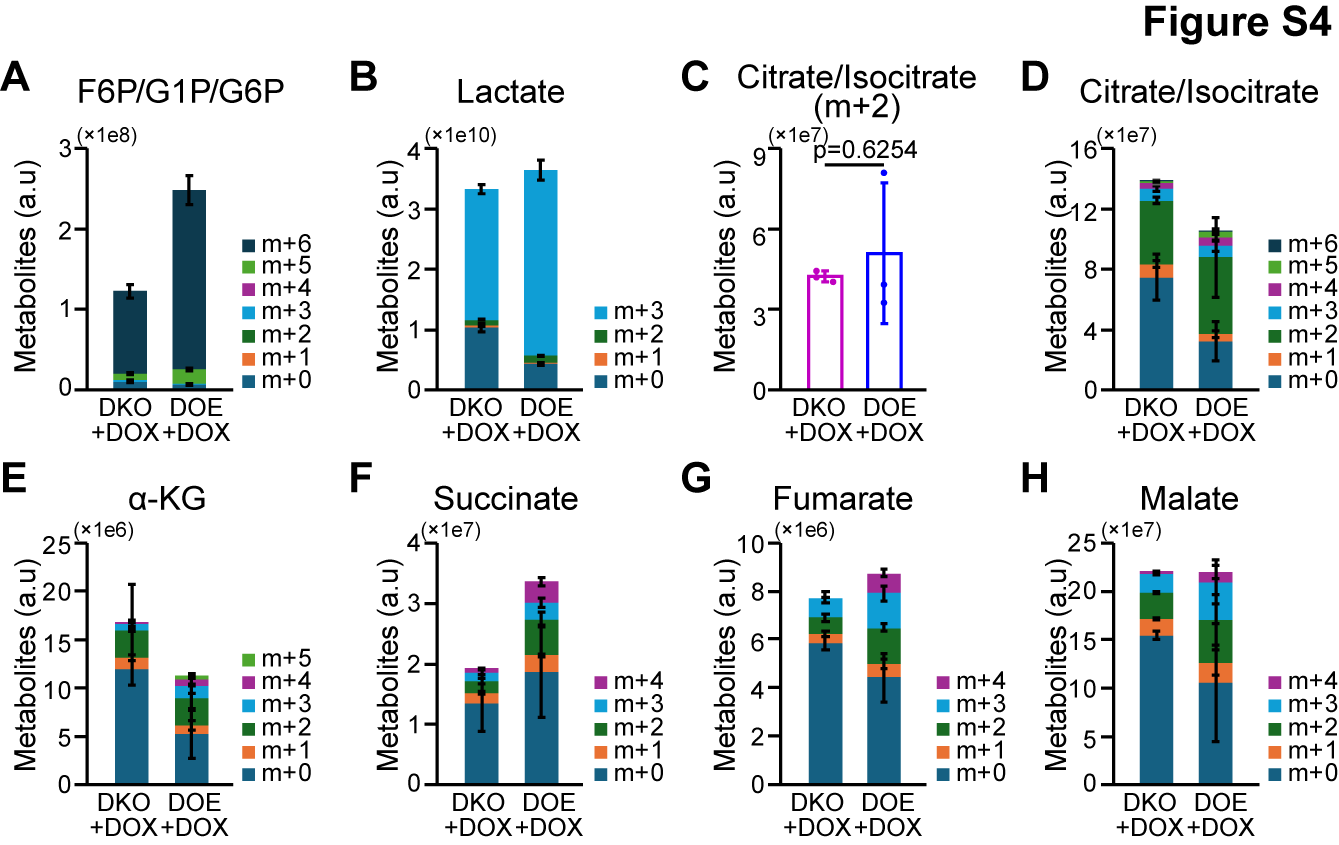

### Figure S5

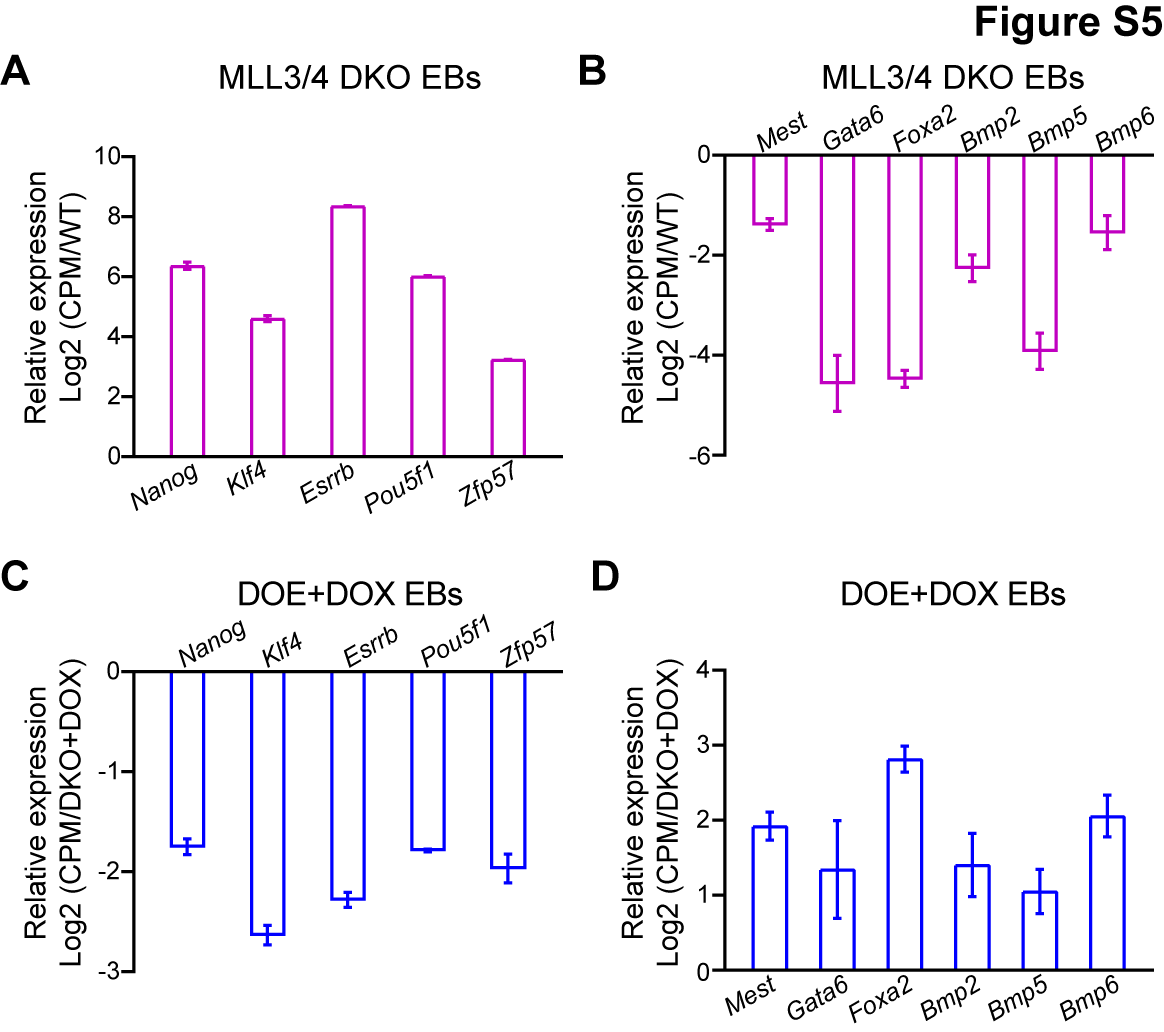
